# Flood-irrigated agriculture mediates climate-induced wetland scarcity for summering sandhill cranes in western North America

**DOI:** 10.1101/2023.11.03.565509

**Authors:** J. Patrick Donnelly, Daniel P. Collins, Jeffrey M. Knetter, James H. Gammonley, Matthew A. Boggie, Blake A. Grisham, M. Cathy Nowak, David E. Naugle

## Abstract

Documenting a species’ extent is often the first step in understanding its ecology and is critical to informing conservation planning. Basic information about species distributions is lacking in many regions of the world, forcing natural resource managers to answer complex ecological questions with incomplete data. Information gaps are compounded by climate change, driving resource bottlenecks that can act as new and powerful demographic constraints on fauna. Here, we reconstructed greater sandhill crane (Antigone canadensis tabida) summering range in western North America using movement data from 120 GPS-tagged individuals to determine how landscape composition shaped their distributions. Landscape variables developed from remotely sensed data were combined with bird locations using cloud computing and machine learning to model distribution probabilities. Additionally, land-use practices and land ownership were summarized within summer range as a measure of use dependence. Wetland variables identified as important predictors of bird distributions were also evaluated in a post hoc analysis using satellite imagery to measure the long-term (1984–2022) effects of climate-driven surface water drying. Wetlands and associated agricultural practices accounted for 1.2% of the summer range but were key predictors of greater sandhill crane occurrence. Bird distributions were patterned primarily by riparian floodplains that concentrated water, wetlands, and flood-irrigated agriculture in otherwise arid and semi-arid landscapes. Findings highlighted the critical role of private lands in greater sandhill crane ecology as they accounted for 78% of predicted distributions. Wetland drying observed in portions of the range from 1984 to 2022 represented an emerging ecological bottleneck that could limit future greater sandhill crane summer range. Study outcomes provide novel insight into the significance of ecosystem services provided by flood-irrigated agriculture that supported nearly 60% of the wetland resources used by birds. Findings suggest greater sandhill cranes function as an umbrella species for agroecology and climate change adaptation strategies seeking to reduce agricultural water use through improved efficiency while also maintaining distinct flood-irrigation practices supporting greater sandhill cranes and other wetland-dependent wildlife. To inform conservation design, we make our wetland and sandhill crane summering distributions publicly available as interactive <u>web-based</u> <u>mapping tools</u>.

## 1 INTRODUCTION

Knowing a species’s geographical extent is often the first step in understanding its ecology and is critical to informing conservation planning (Guisan *et al*., 2013). In many regions of the world, even basic information about species distribution is lacking (Pimm *et al*., 2014), forcing natural resource managers to answer complex ecological questions with incomplete data. Increased frequency and severity of extreme weather and climate events are triggering resource bottlenecks that can act as powerful demographic constraints on fauna and exacerbate other human-induced pressures such as land-use change (Maron, McAlpine and Watson, 2015). Addressing these evolving information gaps requires the integration of GPS animal tracking technologies with satellite imagery and cloud computing to efficiently monitor species’ response to changing ecosystem conditions at regional and continental scales (Pimm *et al*., 2015). Such information can provide the foundation for science-based management necessary for assessing emerging biodiversity risks (Wiens *et al*., 2009).

In western North America, greater sandhill cranes (*Antigone canadensis tabida,* hereafter sandhill cranes) are iconic migratory waterbirds representative of wetland and riparian (hereafter ‘wetland’, Armbruster 1987) ecosystems. Sandhill crane’s annual life cycle (i.e., wintering, migration, and summering) is closely linked to water availability and hydrologic cycles that drive wetland function and irrigated agriculture (Donnelly *et al*., 2021). During summering periods, for example, birds defend traditional breeding territories around wetlands to reduce predator risks by building nest mounds in standing water (Austin, Henry and Ball, 2007). Emergent vegetation at these sites, in turn, supports important food resources for sandhill crane colts, particularly during late summer when seasonal drought limits foraging opportunities in desiccated uplands (Drewien and Bizeau, 1974). Increasing water scarcity driven by warming temperatures and prolonged droughts raises concerns over climate resilience in wetland ecosystems supporting sandhill crane populations. Additionally, reliance on privately owned agricultural lands could expose summering birds to an increased risk of land use change driven by climate adaptation strategies that shift cropping and water usage to practices incompatible with bird needs (Austin, Morrison and Harris, 2018).

Characterizing sandhill crane summering distributions and their relationship to rural private lands agriculture fills a crucial gap in western North America’s climate change adaptation strategies. Recent studies associate broad-scale ecosystem services with riparian floodplains supporting flood-irrigated agriculture that bolster climate resilience through groundwater recharge sustaining instream flows (Kendy and Bredehoeft, 2006; Gordon *et al*., 2020) and coldwater fisheries (Blevins *et al*., 2016). In some regions, these agroecosystems complement broader wetland function, particularly flood-irrigation tied to grass-hay production that supports nearly 60% of temporary wetlands in the Intermountain West, USA (Donnelly et al. *in press*), providing vital resources for migratory waterbirds (Moulton *et al*., 2022). Well-intended efforts to curtail climate change impacts through government programs that increase agricultural water use efficiencies by targeting perceived wasteful flood-irrigation could unintentionally decouple long-standing land-use practices benefiting sandhill cranes and other wetland-dependent wildlife. Documenting sandhill crane summering distributions will provide an important addendum to climate adaptation strategies seeking to reduce overall water consumption on public and private lands through an improved understanding of tradeoffs between changing water management and ecosystem services benefiting these birds.

While regional studies have provided insight into sandhill crane ecology (Littlefield, Stern and Schlorff, 1994; McWethy and Austin, 2009), high dispersal rates and relatively low densities during summering periods have limited our knowledge of landscape drivers structuring bird distributions. To address these information gaps, we marked 120 sandhill cranes with GPS tags and tracked their movements across eight U.S. states and Canadian provinces (Figure 1). A suite of remotely sensed ecological variables representing climate, human disturbance, land cover, and wetlands were then combined with bird locations in a cloud computing platform to model summer-range sandhill crane distribution probabilities. Wetland variables identified as important predictors of bird distributions were evaluated in a post hoc analysis using satellite imagery to measure the long-term (1984–2022) effects of climate-driven surface water drying (Donnelly *et al*., 2020). Agricultural irrigation practices and land ownership were summarized within core sandhill crane summering areas as a measure of land-use dependence. Study outcomes provide novel insight into the significance of wetland scarcity and flood-irrigated agriculture in structuring sandhill crane summering distributions.

**Figure 1.**
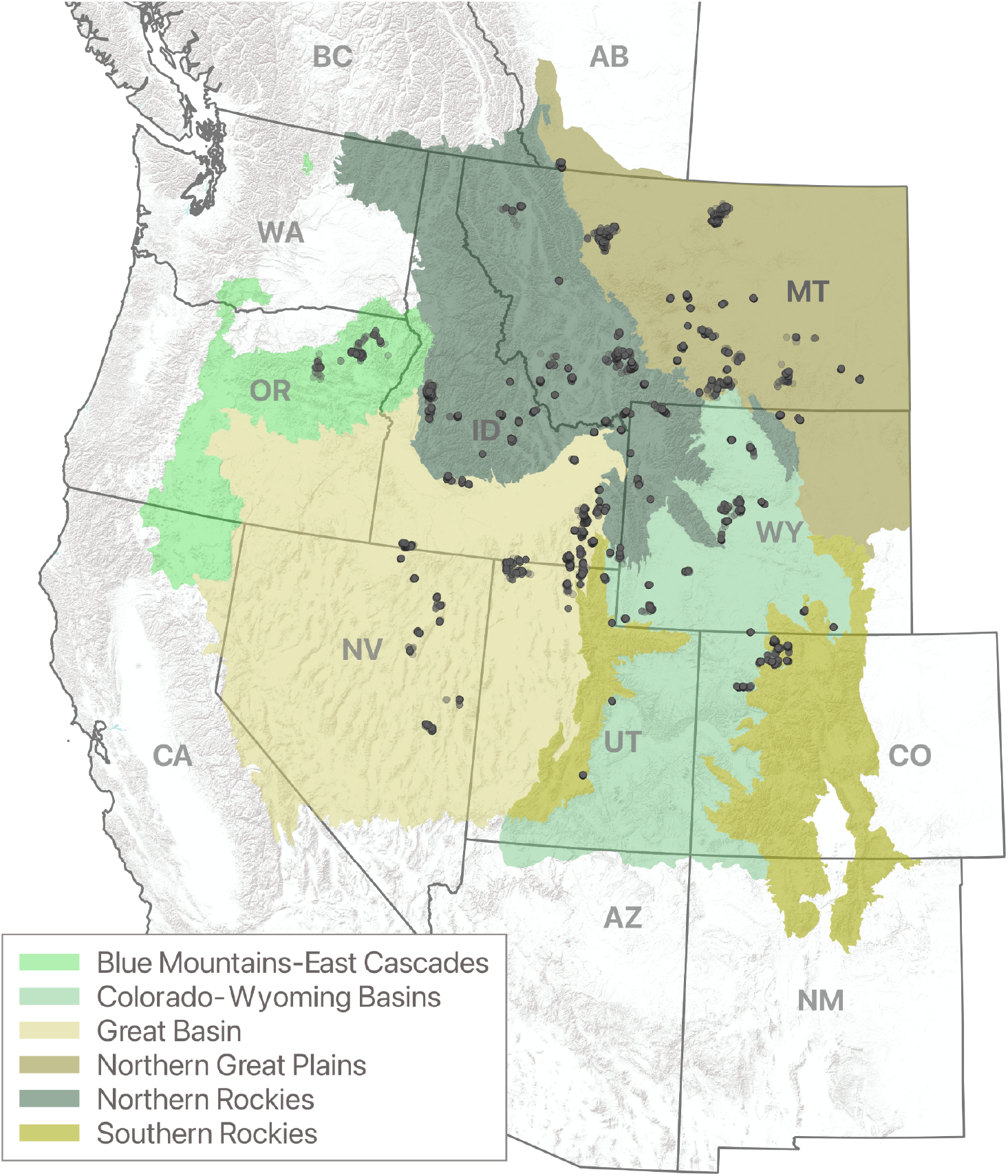
Study area extent defined by modified Level III ecoregions – shown in color. Points identify GPS summering sandhill crane locations collected 2014–22.

## 2 Materials and Methods

### 2.1 Study Area

The study area encompassed portions of the Intermountain West and Northern Great Plains, as defined by North American level three ecoregions (Wiken, Jiménez Nava and Griffith, 2011), including areas overlapping thirteen provinces and states in Canada (Alberta, British Columbia) and the United States (Arizona, California, Colorado, Idaho, Nevada, New Mexico, Montana, Oregon, Utah, Washington, and Wyoming). Ecoregional boundaries included known and suspected sandhill crane summering distributions based on published breeding surveys and GPS location data collected for this study (Drewien and Bizeau, 1974; Littlefield, Stern and Schlorff, 1994; Conring, 2016; Thorpe, Donnelly and Collins, 2022). Individual ecoregions were used to normalize summary results as a means to interpret patterns of land use and wetland change within the birds’ range. Original ecoregional boundaries were largely unmodified but in some instances, were merged or clipped to align with sandhill crane distributions. Ecoregions within the Intermountain West included: 1) Blue Mountains-East Cascades, 2) Colorado-Wyoming Basins, 3) Great Basin, 4) Northern Rockies, and 5) Southern Rockies. The Northern Great Plains encompassed a single ecoregion (Figure 1).

Most of the study area is considered arid to semi-arid, restricting wetlands to 1–3% of the landscape footprint (Tiner, 2003). Water and wetlands are concentrated in valley bottoms along riparian floodplains. In the Northern Great Plains, glaciated pothole wetlands and small (< 0.4 ha) human-constructed ponds for livestock watering are common and can occur in high densities regionally. Riparian wetlands in the Intermountain West are heavily supported by flood-irrigated grass-hay production used for livestock forage production (Donnelly et al. *in press*). The practice diverts and spreads surface water from adjacent streams to irrigate perennial grasses that are seldom tilled and require relatively low chemical inputs (e.g., soil amendments, pesticides, and herbicides) compared to other more industrialized cropping practices. Publicly owned wetlands include lands administered by the U.S. Fish and Wildlife Service (USFWS), Environment Canada, and state wildlife agencies (hereafter ‘wildlife refuges’) managed to support sandhill cranes and other migratory waterbird populations. Other public wetland owners in the United States included the Bureau of Reclamation, the Bureau of Land Management, and the U.S. Forest Service.

The climate is characterized by cold winters and hot/warm summers. Wetland hydrology (i.e., flooding) is induced by local and high-elevation snowmelt and runoff. In the Northern Great Plains, wetland hydrology can also be influenced by spring and summer precipitation. Most wetlands are inundated seasonally from early spring through mid-summer, after which evaporative drying reduces surface water availability.

### 2.2 Crane captures and GPS deployment

Sandhill crane locations used to derive summering distributions were acquired from 120 individual birds captured and fitted with GPS tags. Tag deployments were partitioned among summering (n = 35) and wintering (n = 85) areas (See Appendix A, Figure A1). Breeding status and sex of tagged adult sandhill cranes (n = 112) were unknown but assumed to have minimal influence on our broad-scale assessment of summering distributions. Because sandhill cranes form lifelong pair bonds and maintain close contact with family groups throughout summering, migration, and wintering periods, space-use was considered similar among sexes. Additionally, birds marked as juveniles (n = 8) and identified as non-breeders exhibited movement and space-use patterns similar to other marked birds in the study. Location acquisition rates of individual GPS tags varied from four to 45 points per day. Approximately 685,000 summering bird locations were collected from 2014 to 2022 (Figure 1), with over 85% of days containing seven or more acquisitions per 24 hours. Detailed capture and GPS deployment procedures are provided by Collins et al. (2016) and Boggie et al. (2018).

### 2.3 Defining summering locations

First passage time analysis (FPT) was used as a metric to differentiate sandhill crane summering and migration behaviors (Johnson *et al*., 1992) by identifying spatiotemporal scales that birds interacted with landscapes (Fauchald and Tveraa, 2003). Because FPT is scale-dependent, we calculated sandhill crane movement variance at radii from 1–100 km to distinguish slow localized summering movements from rapid long-distance migration (sensu Le Corre, Dussault and Côté, 2014). We segmented FPT results temporally using Behavioral Change Point Analysis (BCPA; Lavielle, 1999; Lavielle and Teyssière, 2006). This method optimized segmentation of seasonal space-use by minimizing a contrast function (i.e., a function measuring the discrepancy between rapid long-distance migration and an underlying model characterized by slow localized movements). We applied BCPA using a mean contrast function and minimum location use parameter of 10. Differences in GPS acquisition rates among birds did not influence BCPA segmentation due to large-scale migration versus localized movements that were distinguished. Both FPT and BCPA were implemented with R-package adehabitatLT (Calenge, 2011).

Movement segments were classified for individual birds using a rule-based approach linked to seasonal timing and duration of unique space-use patterns. Summering segments were made of bird locations associated with prolonged localized movements beginning in early spring and ending in late summer. Migration segments were classified as rapid long-distance movements occurring on either end of summering periods. All classified BCPA results were exported to a GIS for visual inspection and editing to ensure classifications aligned with observed bird movements. Summering locations were then combined as a representative sample of all individual birds sampled.

### 2.4 Landscape variables

We used a suite of 15 continuous spatially explicit landscape variables to model summering sandhill crane distributions (Table 1). To fill data gaps in available wetland data, we developed novel surface water models to depict the timing and duration of wetland flooding (hereafter “wetland hydroperiod”), a key delimiter of vegetative structure and foraging resources associated with waterbird use (Foti *et al*., 2012). Following methods outlined by Donnelly et al. (2021), we mapped monthly patterns of wetland inundation for all wetland and riparian systems within the study area. Measurements were derived from surface reflectance Landsat 8 and 9 Operational Land Imager satellite imagery using a 30 x 30 meter pixel grid to capture hydroperiod diversity within individual wetlands. Heavily forested wetlands were omitted from our analysis due to ocular masking from tree canopy, making it difficult to measure surface water conditions. Hydroperiods were summarized by totaling the number of months individual pixels were flooded from January to December. Wetlands were then classified as ‘temporary’ (flooded < 2 months), ‘seasonal’ (flooded > 2 and < 9 months), or ‘semi-permanent’ (flooded > 8 months) using standards similar to Cowardin et al. (1979). This process was replicated annually from 2014-2022 to coincide with sandhill crane GPS data collection periods. Large deep water bodies (e.g., reservoirs and large rivers) were omitted from our analysis to remove bias from wetland features that do not directly support sandhill crane summering habitat. A detailed description of methods used to generate wetland hydroperiods is provided in Appendix A.

**Table 1.**
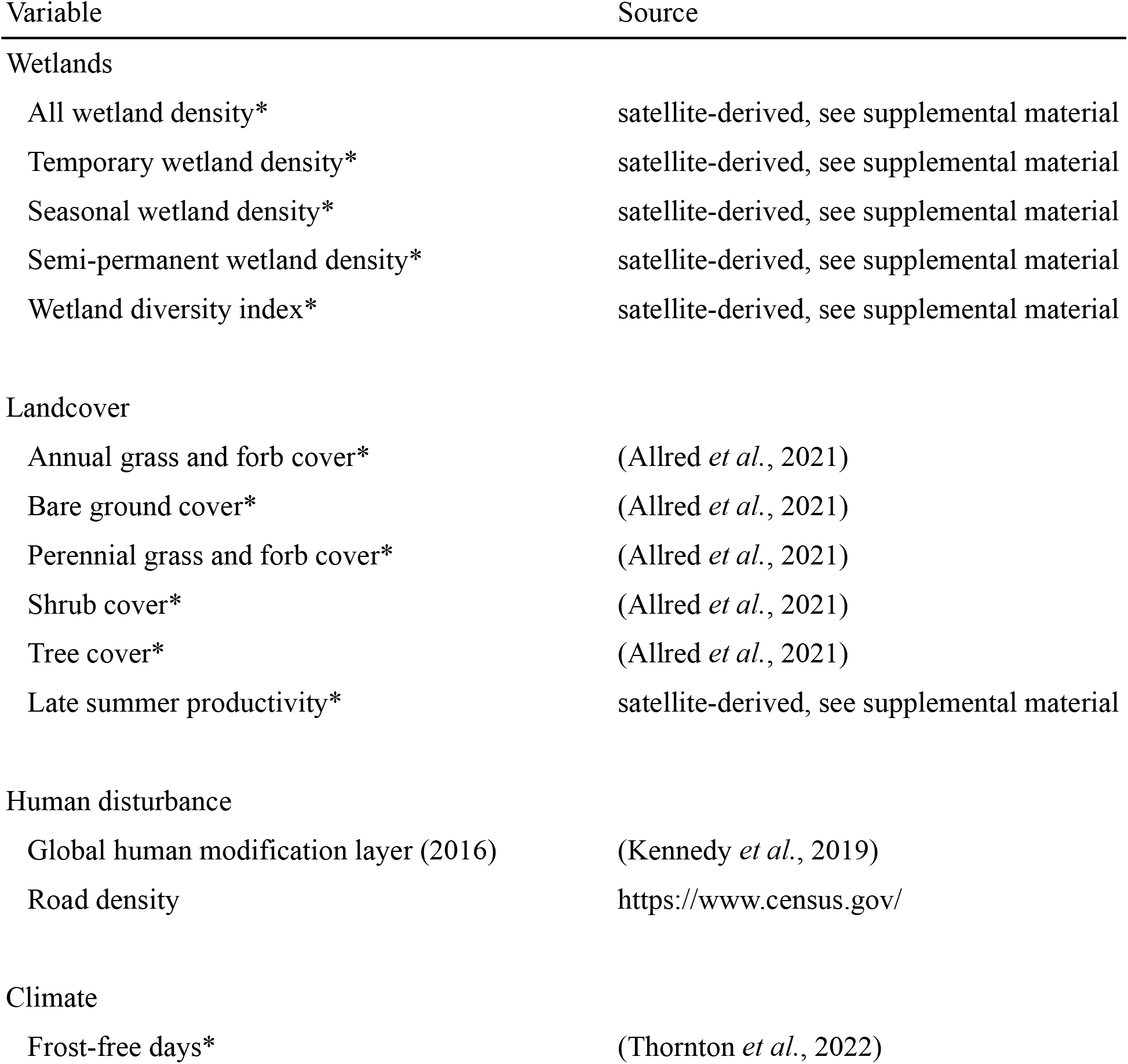
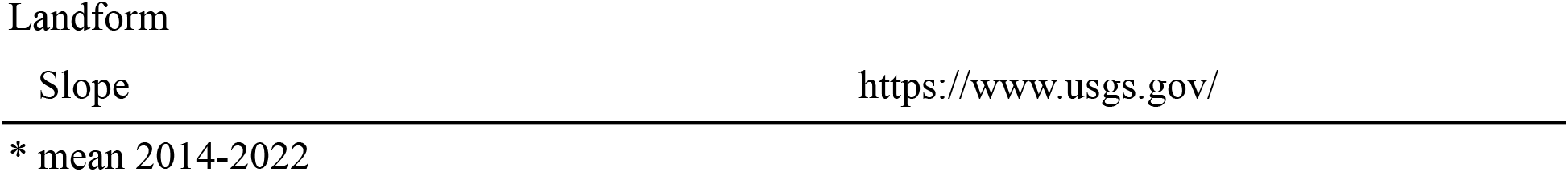
List of variables used to predict sandhill crane summering distributions.

We applied a mean focal filter to wetland distributions to estimate densities within landscapes as a means to capture the gradient of space-use sandhill cranes exhibited around these features. The kernel diameter used for focal calculations (~3 km) was equivalent to the eightieth percentile home range estimates for summering sandhill cranes to ensure outputs encompassed bird movements. Home range estimates (n = 120) were derived for individual birds from 2014–2022 using GPS locations and minimum convex hull calculations. Wetland density measures were replicated for all wetland hydroperiods combined and individually for each hydroperiod class. We accounted for wetland diversity as a factor influencing sandhill crane distributions by applying Simpon’s index of diversity (Simpson, 1949):

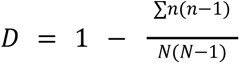

where *n* = the area of each wetland hydroperiod class (temporary, seasonal, or semi-permanent) and *N* = the total number of wetland hydroperiod classes (temporary, seasonal, semi-permanent). A mean focal filter approach previously described was used to estimate wetland diversity within the median home range of summering sandhill cranes.

All wetland hydroperiod and wetland diversity measures presented were calculated as an average during nesting and early colt-rearing periods (April 1 to June 30) for each year GPS locations were acquired (2014–2022). Results were linked with annual bird locations to align resource conditions with space use over time.

### 2.5 Crane distribution modeling

We modeled the relationship between sandhill crane GPS locations and environmental factors to predict summering distributions using a Random Forest regression algorithm (Breiman, 2001). Random Forest uses machine learning to produce predictive models that account for nonparametric and highly complex ecological interactions (Mi *et al*., 2017). This modeling approach is less sensitive to collinearity issues among predictors while remaining robust to overfitting (Culter *et al*., 2007). Hyperparameter tuning was used to identify the optimal number of trees needed (n ~ 4000) to maximize model accuracy (Oshiro, Perez and Baranauskas, 2012). Because sandhill cranes exhibited constrained movement patterns with clustered space-use in small areas due to nesting and early colt-rearing behaviors, locations were sampled to remove bias from spatial autocorrelation when formatting Random Forest training data. A total of 20,000 locations from 685,000 GPS points collected were selected randomly using a rule-based process that restricted the selection of locations within a distance of 400m from other selected points. Pseudo-absence locations needed for model training were generated within the study area and combined with sampled sandhill crane GPS locations (n = 20,000). We optimized Random Forest performance that can be sensitive to uneven sampling among classes by maintaining a balanced ratio of presence-absence locations (Liu, Newell and White, 2019).

To reduce the potential of false absences, GPS sandhill crane locations were buffered by 1.5 km to exclude pseudo-absence points from sandhill crane use areas (VanDerWal *et al*., 2009). Buffered distance approximated the size of the median sandhill crane summer home range area, as described previously. The rfUtilites package in R (Evans and Murphy, 2015) was used to test and remove multicollinear variables. All variables were retained. To assess variable importance in Random Forest, we used the sum of squared errors measured as the total increase in node purity resulting from variable splitting averaged over all trees in the model. Partial dependence plots for variables were developed as a graphical depiction of the marginal effects influencing sandhill crane resource selection propensity.

A holdout ratio of 20% (n ~ 8,000) was randomly selected to evaluate model performance using k-fold (k = 5) cross-validation and a classification probability threshold of 0.65. Model dispersal constraints were implemented through expert opinion conducted by regional state and provincial wildlife biologists (n > 25) familiar with local sandhill crane distributions (Hengeveld and Haeck, 1982). Feedback from this process was used to inform model parameters and to remove areas of unoccupied habitat on the fringe of the bird’s predicted summer range. All model results were interpreted using a core area approach that summarized bird use and landscape variables within predicted high probability (≥0.65) use areas (hereafter “core areas”).

### 2.6 Wetland trends

High-ranking wetland variables identified by the Random Forest model were applied to a post hoc analysis using Landsat satellite imagery to measure long-term (1984–2022) wetland hydrology in sandhill crane core summering areas. Following methods outlined by Donnelly (2020), surface water patterns were monitored annually as three-month means that coincide with sandhill crane nesting and early colt-rearing (Apr-Jun). Increased surface water area during this time correlates positively to sandhill crane nest success (Austin, Henry and Ball, 2007). Analyses were binned by hydroperiod (i.e., temporary, seasonal, and semi-permanent) to examine change within wetland factors used to predict sandhill crane distributions. Wetland hydroperiod classes were modeled using mean conditions during the sandhill crane GPS data collection period (2014–2022) and were held constant over time. Trends were measured by fitting Kendall’s Tau-b rank correlation to pixel surface water change through time. Trend measurements were partitioned eco-regionally to isolate spatial differences in hydrologic function (Lauenroth, Schlaepfer and Bradford, 2014).

### 2.7 Land-use

Core areas were summarized by private, public, and tribal land management designations to measure crane land-use reliance. Private land-use was categorized as 1) lands protected under conservation easement, 2) flood-irrigated grass-hay cultivation (hereafter “grass-hay”), or 3) other private lands. Public lands were partitioned by agencies (e.g., Bureau of Land Management, Forest Service). Tribal lands were denoted as 1) grass-hay or 2) other tribal lands. Public and protected lands were defined using the Protected Area Database for the United States and the Canada Lands Digital Cadastral Dataset. Grass-hay was defined using Donnelly et al.’s (*in press*) grass-hay production layer. This dataset was chosen because of the strong spatial correlation between grass-hay delineations and riparian floodplains 一 known to support flood-irrigated wetlands (sensu Gordon *et al*., 2020) and summering bird distributions in portions of their range (McWethy and Austin, 2009). Because wetland hydrology was identified as a key component in structuring sandhill crane summering distributions, wetland hydroperiods were also summarized by private, public, and tribal land management designations.

### 2.8 Data Processing

All image processing and raster-based analyses were conducted using Google Earth Engine, a cloud-based geospatial processing platform (Gorelick *et al*., 2017). All GIS analyses were performed using QGIS (QGIS Development Team, 2020). Plotting and statistical analyses were generated using the R environment (R Core Team, 2019; RStudio Team, 2019), including R-packages not previously mentioned; randomForest (Liaw, Wiener and Others, 2002), and Tidyverse (Wickham *et al*., 2019).

## 3 RESULTS

### 3.1 Summering distributions

Core sandhill crane summering areas accounted for ~5% (7.6 million ha) of the total estimated range (~145 million hectares) in western North America, encompassing eleven states and provinces in the U.S. and Canada (Figure 2). Eastern portions of the Great Plains ecoregion and southern portions of the Colorado-Wyoming Basins, Great Basin, and Southern Rockies ecoregions were modified to represent occupied sandhill crane range. Northern areas of the Northern Rockies ecoregion were also modified (Figure 2). The percentage of core area was greatest in the Northern Great Plains, containing 34.5% of the total core area, followed by the Northern Rockies (19.1%), Great Basin (15.7%), Colorado-Wyoming Basins (14.8%), Southern Rockies (8.0%), and Blue Mountain-East Cascades (7.8%) ecoregions. By state, Montana accounted for 41.6% of core area, followed by Wyoming (16.3%), Idaho (12.8%), Oregon (7.9%), Utah (7.4%), Colorado (7.0%), Nevada (2.9%), California (2.6%), and Washington (0.2%). Canada comprised less than 2.0% of the core area distribution located in southwest Alberta and southeast British Columbia. The model mean square error for k-fold cross-validation (k = 5) was 0.29.

**Figure 2.**
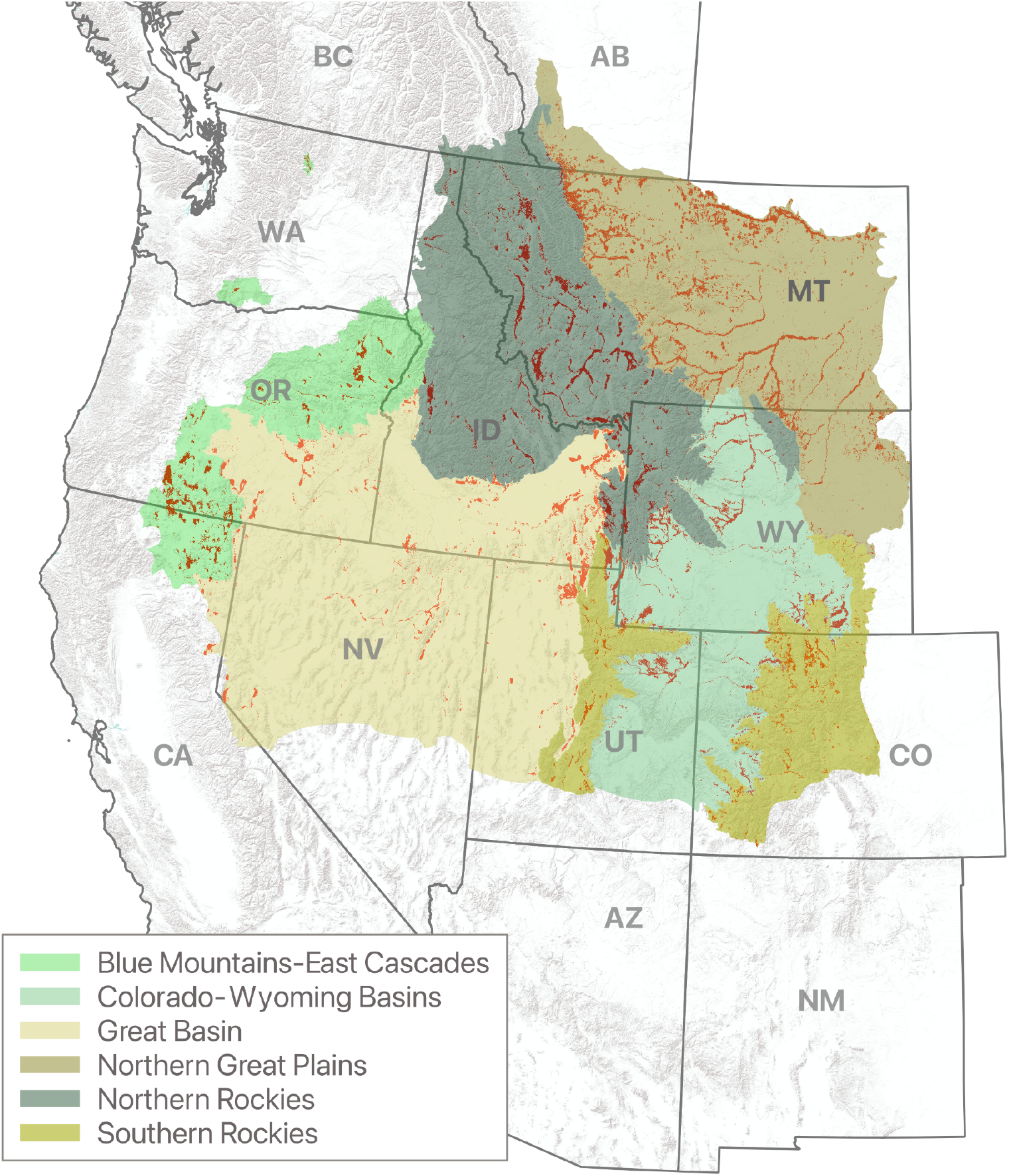
Sandhill crane core summering distributions shown in red as occurrence probabilities ≥ 65%. Modified ecoregional polygons encompass estimated summer range extent.

### 3.2 Landscape variables

Sandhill crane summering distributions were structured primarily by wetland density, wetland diversity, and slope (Figure 3). Resource selection propensity for temporary wetland density, overall wetland density, wetland diversity, and seasonal wetland density increased sharply at measures above zero and was maintained as measures increased (Figure 4). Sandhill crane occurrence probabilities were negatively correlated to slope and declined to near zero in areas greater than 20%. Other high-ranking variables included semi-permanent wetland density and late summer vegetative productivity. Bird probabilities were positively correlated to semi-permanent wetlands at low densities and negatively correlated to those > 30m^2^/km^2^. Sandhill crane relationships to late summer vegetative productivity were bimodal, showing high correlations to NDVI values < 0 and > 0.2. This pattern was likely caused by bird use of shallow temporary and seasonal wetlands that typically have negative NDVI values due to the absorption of near-infrared wavelengths by water. Human disturbance measures (i.e., road density and global human footprint) were negatively correlated to sandhill crane occurrence probabilities but were not as important to shaping bird distributions when compared to other variables. Sandhill crane summering areas were generally associated with areas between 125 to 250 frost-free days. Tree cover and bare ground were negatively correlated with occurrence. Shrub, annual grass, and forb cover were least important to structuring sandhill crane summering distributions.

**Figure 3.**
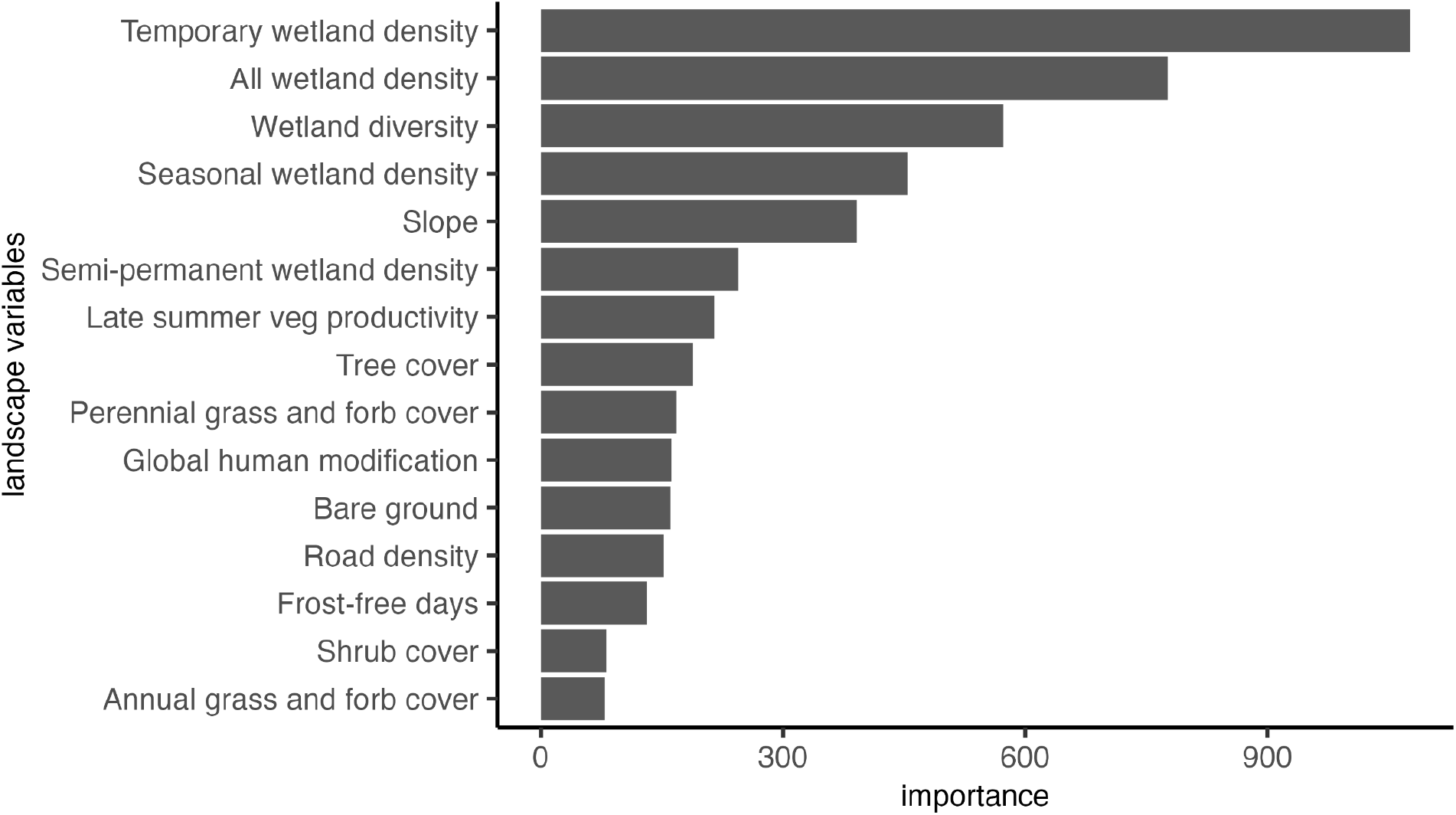
Random Forest variable importance for sandhill crane distribution predictions measured as the total increase in node purity resulting from variable splitting, averaged over all trees.

**Figure 4.**
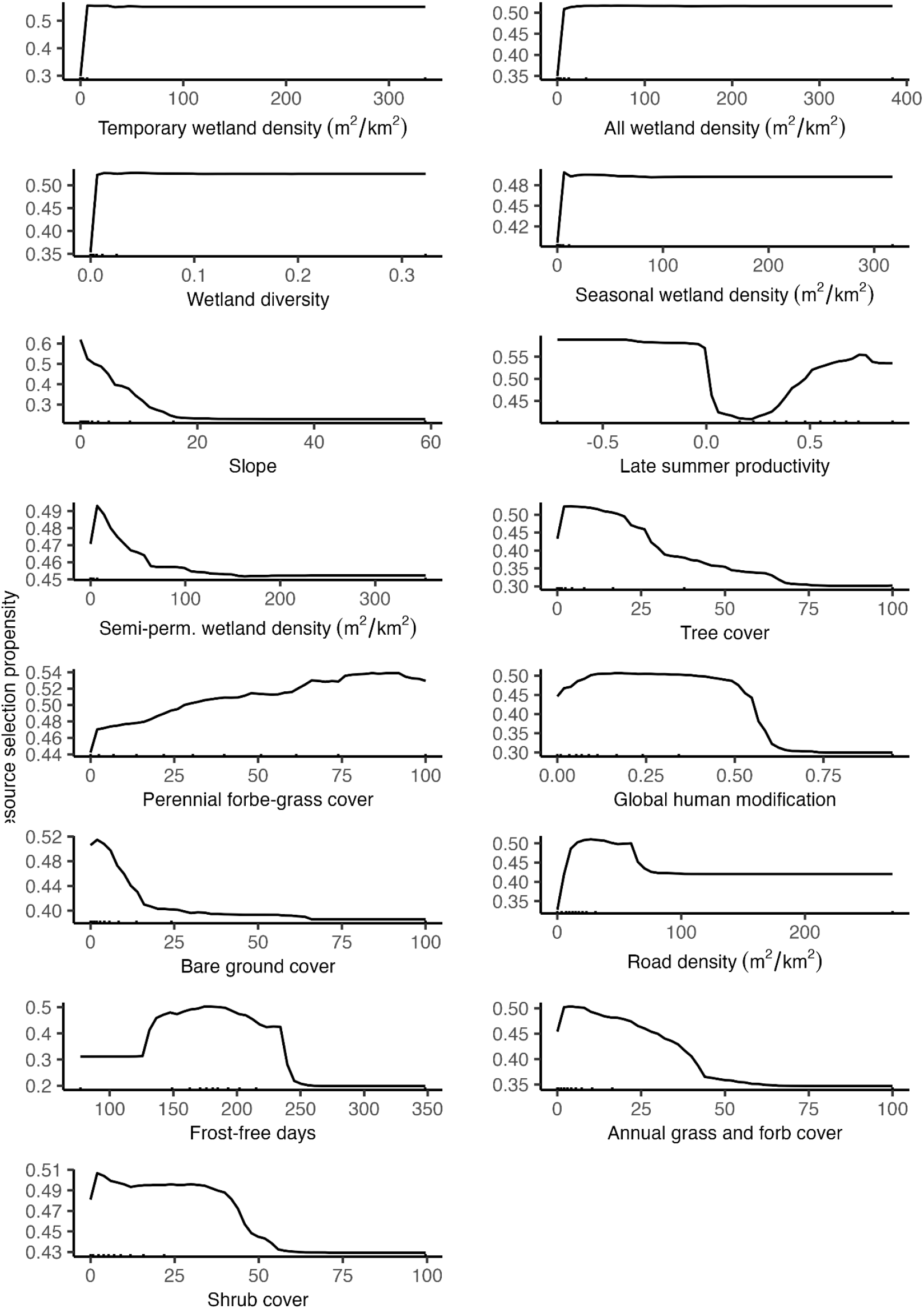
Partial dependence plots depicting marginal effects as a measure of resource selection propensity for each landscape variable used in Random Forest model predictions of sandhill crane summering distributions.

### 3.3 Land ownership

Only 40.3% of sandhill crane summer range was privately owned but it accounted for approximately 78% of the range in core summering areas (Figure 5). In the Intermountain West, core summering areas overlapped 93% of grass-hay agricultural practices. Public lands accounted for approximately 16% of core summering areas, most of which were administered by the U.S. Forest Service, the Bureau of Land Management, and the U.S. Fish and Wildlife Service (see Appendix B, Table B1). Tribal nations, including areas of Tribal grass-hay cultivation, accounted for approximately 6% of core summering areas, primarily on the lands of Blackfeet, Shoshone-Paiute, Eastern Shoshone, and Northern Arapaho peoples (Figure 5).

**Figure 5.**
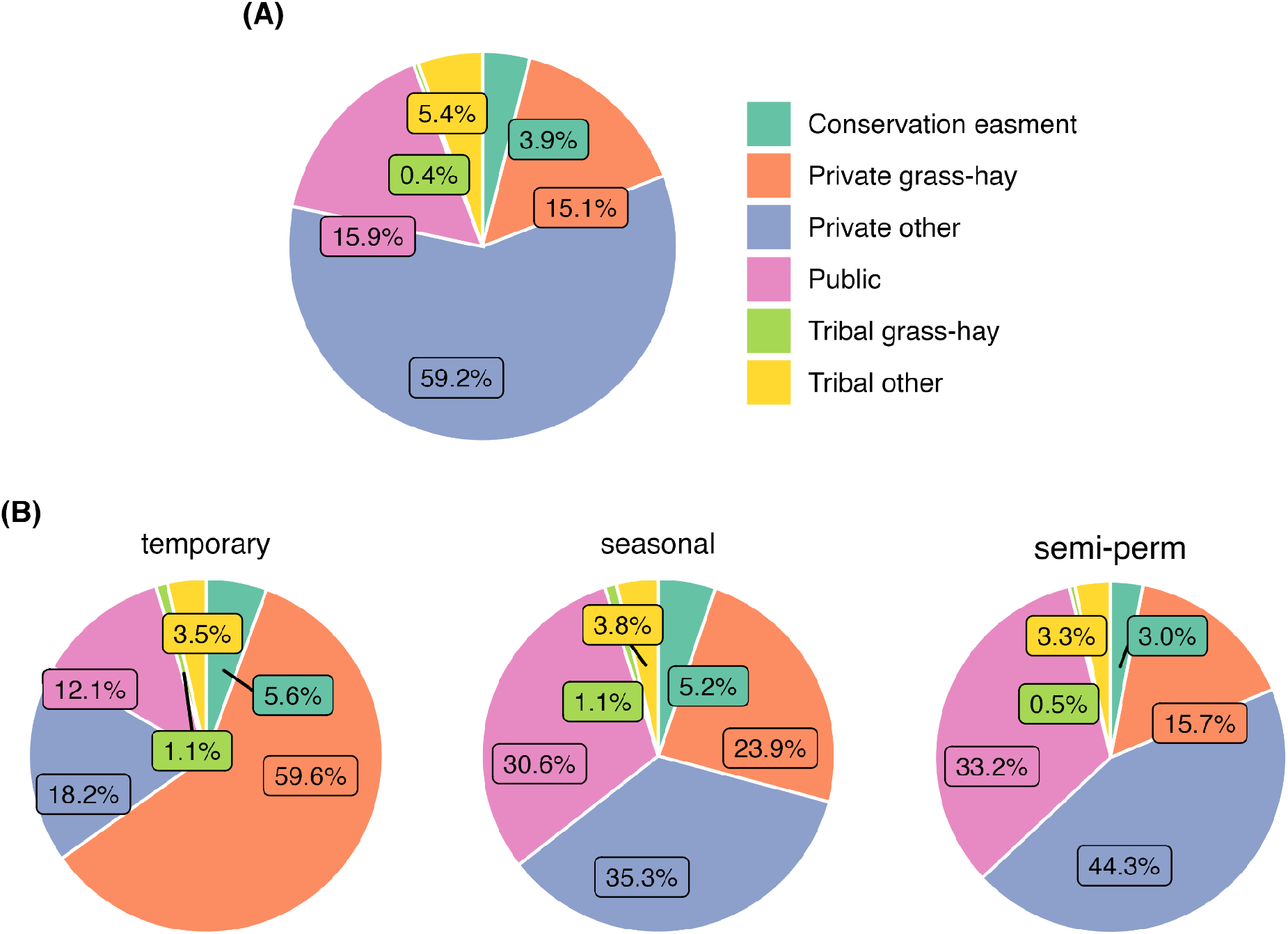
Proportional land ownership, land-use (A), and wetland distributions (B) in sandhill crane core summering areas. Wetlands are summarized by hydroperiod classes (temporary, seasonal, and semi-permanent).

Ownership and land-use practices associated with core area wetland features differed from overall ownership patterns (Figure 5). Core areas encompassed 733,123 ha, 574,654 ha, and 330,512 ha of flooded temporary, seasonal, and semi-permanent wetlands. Privately owned grass-hay production supported 59.6% of temporary wetlands, while an additional 18.2% and 12.1% were attributed to other private and public lands. Seasonal wetlands were distributed relatively evenly between public, private, and grass-hay lands, accounting for 30.6%, 35.3%, and 23.9% of abundance. Private and public lands accounted for over three-quarters of semi-permanent wetlands, representing 44.3% and 33.2% of abundance while grass-hay production on private lands accounted for 15.7% of the total. The U.S. Fish and Wildlife Service, U.S. Forest Service, State Wildlife Agencies, and the Bureau of Land Management accounted for most publicly owned temporary, seasonal, and semi-permanent wetlands (see Appendix B, Table B2-4). Combined Tribal lands (including associated grass-hay) and conservation easements accounted for approximately one to five percent of temporary, seasonal, and semi-permanent wetlands within core areas.

### 3.4 Wetland trend

Wetlands accounted for only for 1.2% of sandhill crane summer range. Wetland surface water monitoring in core areas during sandhill nesting and early colt-rearing periods (April 1 to June 30) showed drying trends correlated to hydroperiod ephemerality. Areas of negative Kendall’s Tau-b rank correlation (i.e., reduced duration or frequency of flooding) were greatest in temporary wetlands and increasingly less pronounced in wetlands characterized by seasonal and semi-permanent hydrologies (Figure 6). Overall Kendall’s Tau-b summaries showed a majority of temporary and seasonal wetlands trending towards dryer states. Rates of change within ecoregions were variable with drying most pronounced in the Great Basin, which showed 71.3% and 65.5% of temporary and seasonal wetlands trending toward dryer states since 1984 (Table 1). The Northern Rockies, Southern Rockies, and Blue Mountain-East Cascades ecoregions also experienced drying in over 50% of temporary wetland areas, indicating long-term declines in hydrologic function. The Colorado-Wyoming Basins and the Northern Great Plains showed stable to wetter trends across all wetland hydrologies. Semi-permanent wetlands within core areas remained relatively stable across all ecoregions, with most wetlands exhibiting Kendall’s Tau-b measures ≥ 0. See Appendix B, Figures B2-7 for Kendall’s Tau-b rank correlation distributions by ecoregion.

**Figure 6.**
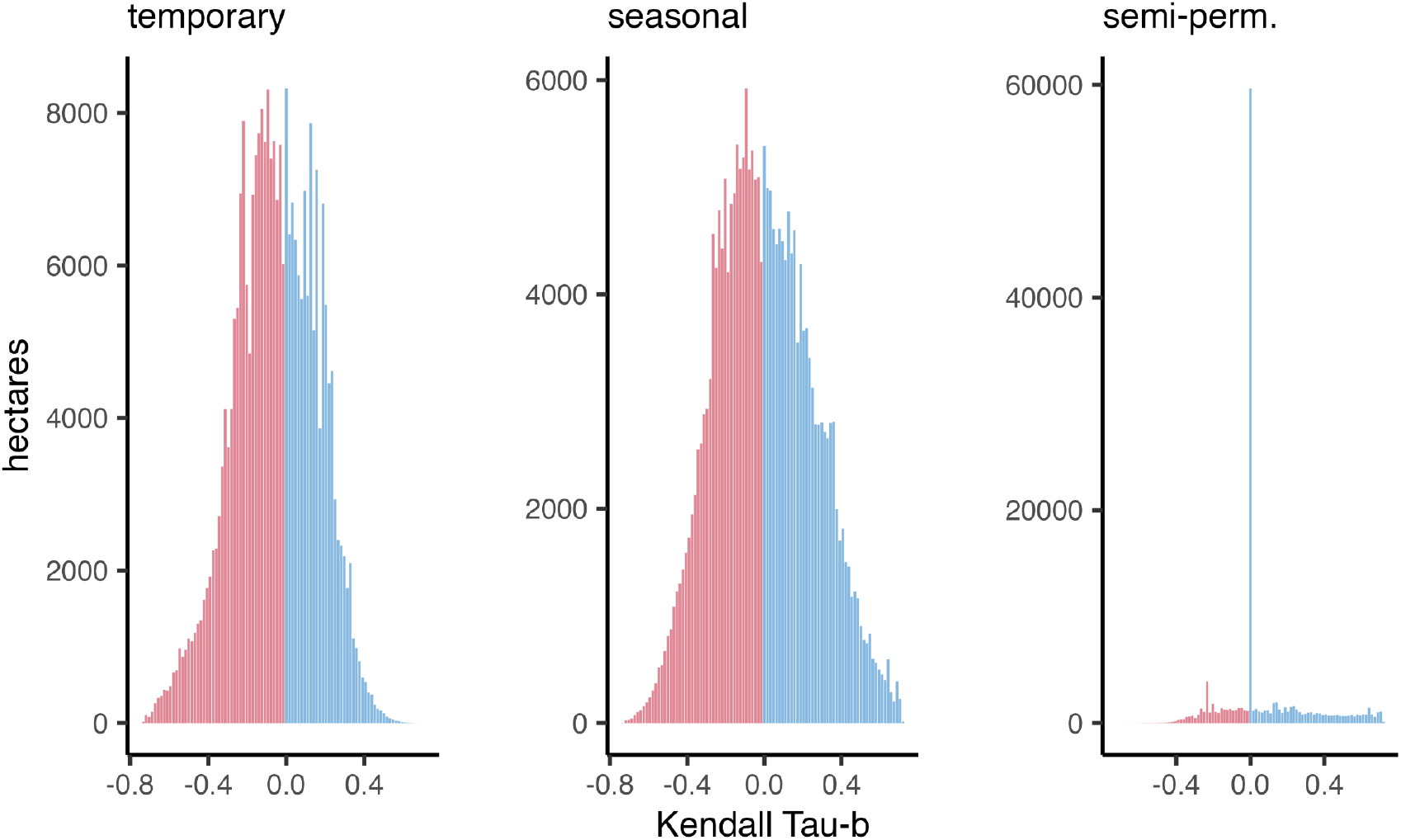
Kendall’s Tau-b rank correlation distributions by wetland hydroperiod class. Summaries represent all wetlands within sandhill crane core areas. Negative values indicate wetland drying associated with reduced duration or frequency of flooding since 1984.

**Table 1.** Proportion of wetland area drying within sandhill crane core areas as determined by negative Kendall’s Tau-b rank correlation values. Summaries are partitioned by ecoregion and wetland hydroperiod. Wetland conditions were measured for each year from 1984–2022 during sandhill crane sandhill nesting and early colt-rearing periods (April 1 to June 30).

|  | temporary | drying | %drying | seasonal | drying | %dryng | semi-perm | drying | %drying |
| --- | --- | --- | --- | --- | --- | --- | --- | --- | --- |
| Blue Mountains - East Cascades | 26,966 | 16,004 | 59.3% | 49,332 | 24,963 | 50.6% | 22,708 | 7,642 | 33.7% |
| Colorado-Wyoming Basins | 86,003 | 38,883 | 45.2% | 38,662 | 13,380 | 34.6% | 36,717 | 2,653 | 7.2% |
| Great Basin | 68,296 | 48,678 | 71.3% | 57,601 | 37,729 | 65.5% | 39,087 | 9,545 | 24.4% |
| Northern Great Plains | 32,423 | 15,901 | 49.0% | 40,239 | 13,464 | 33.5% | 55,136 | 6,451 | 11.7% |
| Northern Rockies | 50,978 | 34,808 | 68.3% | 52,951 | 21,520 | 40.6% | 30,510 | 2,021 | 6.6% |
| Southern Rockies | 21,537 | 12,921 | 60.0% | 11,183 | 4,334 | 38.8% | 6,697 | 405 | 6.0% |

## 4 DISCUSSION

Our analysis is the first to estimate sandhill crane summer range in the American West. The results of this study fill an essential gap in our understanding of bird distributions and the role wetland scarcity and agriculture play in structuring sandhill crane occurrence. Although wetlands accounted for 1.2% of their summer range, they were key predictors of occurrence probabilities. Core areas were structured primarily by temporary wetlands supported by grass-hay agriculture concentrated in riparian floodplains (Donnelly et al. in press). Wetland drying observed in core areas from 1984 to 2022 was consistent with climate-induced trends observed in regional studies (Donnelly *et al*., 2020, 2022), representing an emerging ecological bottleneck that could limit sandhill crane summering range in the future.

Our work indicates a significant sandhill crane range expansion into the Northern Great Plains. Drewien and Bizeau’s (1974) original range estimations documented historical eastward movement in 1972 with the furthest east-known breeding pair in west-central Wyoming near Ocean Lake. Over the past 50 years, we show core summering area expansion of 350 km east and 650 km north, encompassing nearly the entire rangeland ecosystem east of the Rocky Mountains in Montana and Wyoming, including portions of Alberta, Canada. Annual surveys conducted from 1996 to present identify moderate long-term population growth that is attributed primarily to a 240% and 210% increase in sandhill crane abundance in Montana and Wyoming (Thorpe, Donnelly and Collins, 2022), affirming Northern Great Plains range expansion identified by our analysis. Hunting pressure targeting lesser sandhill cranes *(Antigone canadensis canadensis)* in the mid-continent population may act as a potential barrier to further eastward expansion due to incidental harvest, preventing the establishment of breeding pairs (Krapu and Brandt, 2010). Despite a gap in sandhill crane location data in some western portions of the summering range (i.e., California, Nevada, Oregon, and Washington), our core area model aligned well with regional surveys conducted in the mid-1990s by Littlefield et al. (1994) and Rawling (1992), as confirmed by our independent review from regional wildlife managers. Additionally, our results suggest that outside the Great Plains, bird distributions have remained relatively stable within their previously known summer range (Drewien and Bizeau, 1974; Rawling, 1992; Littlefield, Stern and Schlorff, 1994).

At broad scales, wetland density and diversity were the primary factors underpinning core area distributions, partly dismissing the notion that human disturbance acts as a mechanism to limit occurrence during the breeding period (Drewien and Bizeau, 1974; Armbruster, 1987). Low variable importance scores for ‘road density’ and global ‘human disturbance’ variables were reflected in bird movements that showed individuals consistently occupying space adjacent to human dwellings and road networks. Early observations describing sandhill crane habitat suitability were confined mainly to National Wildlife Refuges that provided birds with isolated nesting opportunities and may have biased interpretations of their need for seclusion during breeding (Drewien and Bizeau, 1974; Armbruster, 1987). Sandhill crane range expansion identified here may be an indicator of changing density dependence in core summering areas that has reduced the availability of isolated breeding locations, forcing birds to rely on alternative resources. Because this study did not examine sandhill crane vital rates, it was unclear if bird tolerance for human disturbance functions as a biological sink, where adaptive plasticity for marginal environmental conditions results in reduced nest success and colt survival. Slow life history traits relative to other waterbirds (i.e. low annual fecundity and high adult survival) in sandhill cranes may also mask detectability of these behavioral and habitat interactions at the population level (Drewien, Brown and Kendall, 1995).

Our model depictions of declining surface water likely reflect a new normal in wetland conditions that could impact resource availability in core sandhill crane summer range, particularly in the Intermountain West’s Great Basin and Northern Rockies ecoregions where over two-thirds of temporary wetlands experienced long-term drying. Wetlands play a crucial role in sandhill crane nesting ecology as evidenced by their attraction to water and its positive role in nest survival (McWethy and Austin, 2009). Corroborating our evidence of wetland scarcity for future nesting populations are climate projections showing a 1–3°C increase in Great Basin summer temperatures by 2020–2050 (Snyder *et al*., 2019). Increasing temperatures across the Intermountain West are predicted to decrease snowpack runoff supporting wetland hydrology while simultaneously increasing agricultural water needs through elevated evaporative demand from crops (Mix, Rast and Lopes, 2009; Elliott *et al*., 2014).

Sandhill cranes’ symbiotic relationship with private lands agriculture is foundational to structuring this species’ summer distribution, as evidenced by core summering areas overlaying 93% of flood-irrigated grass-hay production in the Intermountain West. Our results suggest increasing water scarcity due to warming temperatures and prolonged drought events are the primary threat to maintaining flood-irrigation practices supporting summering birds. Growing urban water demands may also compound projected climate shortfalls by negatively impacting irrigated agriculture in some core areas (Schaible and Aillery, 2017). Efforts in Colorado and Nevada, for example, have proposed agricultural fallowing through the purchase and repurposing of rural irrigation water for municipal use (Thorvaldson and Pritchett, 2006; Welsh and Endter-Wada, 2017). Such scenarios frequently require out-of-basin water transfers, reducing local wetland availability supported through irrigation while eliminating local ecosystem services benefiting climate resilience in riparian ecosystems (Blevins *et al*., 2016; Zhuang, 2016). Additionally, loss of irrigation can increase subdivision risk that removes wildlife-compatible land-use practices in rural landscapes as producers sell off land for development due to its reduced agricultural value (Dozier *et al*., 2017).

In the Northern Great Plains, wetland resilience and sandhill crane range expansion may in part be an ecosystem service provided by agriculture. Ketchum et al. (*in press*) found irrigated agriculture in the region is primarily confined to river floodplains promoting groundwater recharge by growing crops over highly permeable soils allowing the return of unconsumed water to the system. We highlight similar ecosystem services tied to riparian (flood-irrigated) grass-hay production in the Intermountain West (Kendy and Bredehoeft, 2006; Gordon *et al*., 2020); however, this practice represents less than three percent of irrigated lands regionally (Donnelly et al. *in press*), the remainder of which is supported by out-of-floodplain surface water diversions or groundwater pumping that may have limited groundwater recharge or return flow benefits. Additionally, high densities of small (< 0.4 ha) ponds constructed for livestock watering, evident in our wetland mapping data (Figure 7), may have offset patterns of landscape drying and increased surface water availability in rangeland ecosystems that previously lacked natural wetlands to sustain summering crane populations. While our analysis does not directly quantify stock pond abundance, their documented proliferation in the Great Plains region has been identified as a driver of increased biodiversity in agricultural landscapes (Swartz and Miller, 2021).

**Figure 7.**
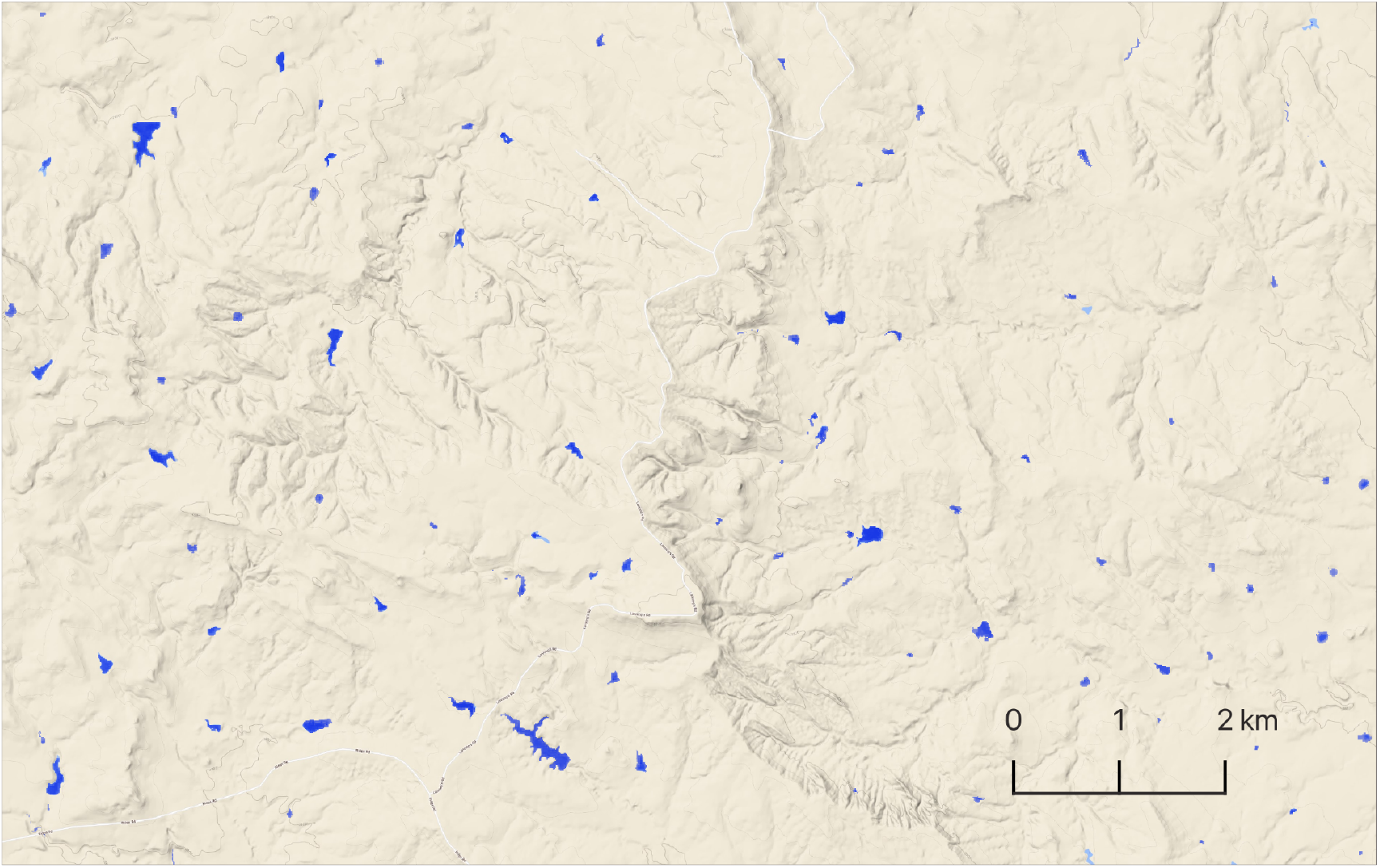
Stock ponds density example from Northern Great Plains, Southeast Montana. Surface water shown in blue.

Tribal lands accounted for a relatively small proportion, approximately six percent, of sandhill crane core area but included sites of relatively high ecological value for migratory waterbirds. Core area distribution on the Fort Hall Reservation lands of the Shoshone-Bannock Tribes (Great Basin ecoregion), for example, are also recognized as a regionally significant shorebird migration site supporting 50,000-80,000 individuals and thirty species annually (Senner, Andres and Gates, 2016). Likewise, core area distribution on the Wind River Reservation lands of the Eastern Shoshone and the Northern Arapaho Tribes (Colorado-Wyoming Basins ecoregion) also support important spring and fall migration staging locations for sandhill cranes summering across the eastern half of their range (Donnelly *et al*., 2021). Senior water rights often associated with tribal lands (Sanchez, Edwards and Leonard, 2023) may elevate their importance to sandhill cranes and other wetland-dependent wildlife as projected climate scenarios indicate increasing water scarcity across most of the summering range. Increasing Tribal engagement can also provide opportunities to integrate traditional ecological knowledge into broader conservation strategies that may benefit from the perspective of indigenous stewardship (Leonetti, 2010).

From a broader ecosystem perspective, our findings suggest sandhill cranes function as a unique umbrella species for agroecology and climate change adaptation strategies considerate of wetland-dependent wildlife and solutions for improved water security. Intrinsic linkages between wetlands and agricultural ecosystem services, represented by our bird distributions in the Intermountain West and Northern Great Plains, support a spatially explicit conservation framework capable of targeting maintenance of beneficial flood-irrigation practices (i.e., grass-hay) while implementing adaptive measures for out-of-floodplain cultivation that reduces overall water use through maximized efficiency. Moreover, sandhill cranes have a long heritage of cultural values in western North America, supporting regional festivals celebrating their seasonal arrivals that have become important drivers in rural economies ーmaking these birds ideal ambassadors for sustainable wetland ecosystems in agricultural landscapes. The cultural significance of sandhill cranes extends to many Native American Tribes that have revered these birds for centuries (Johnsgard, 2017). To inform conservation design, we make our wetland and sandhill crane core summering data publicly available as interactive <u>web-based mapping tools</u>. We encourage the use of our results to inform conservation solutions through collaborative and proactive decision-making among local and regional stakeholders while considering the social, ecological, and economic factors of conservation and management actions at local scales.

## Supporting information

AppendixA

AppendixB

## ACKNOWLEDGEMENTS

We thank the late John P. Taylor, (frm) Senior Wildlife Biologist, Bosque del Apache National Wildlife Refuge, for his ongoing inspiration and original vision that helped to make this work a reality. We thank Idaho Department of Fish and Game, U.S. Fish and Wildlife Service - Division of Migratory Bird Management, and Davis College of Agricultural Sciences and Natural Resources for funding that made this work possible. We recognize Colorado Parks and Wildlife, Idaho Department of Fish and Game, Oregon Department Fish and Wildlife, Montana Fish Wildlife and Parks, New Mexico Game and Fish and the U.S. Fish and Wildlife Service Migratory Bird Program - Southwest Region for their support and coordination of sandhill crane capture and GPS deployment. Views in this manuscript from United States Fish and Wildlife Service authors are their own and do not necessarily represent the views of the agency. Any use of trade, firm, or product names is for descriptive purposes only and does not imply endorsement by the U.S. Government.

## DATA ACCESSIBILITY

Data visualization and interactive mapping tools are available here.

## AUTHORS’ CONTRIBUTIONS

J.P.D. and D.P.C. contributed toward concept and design; J.P.D. analyzed the data; D.P.C., J.H.G., J.M.K., B.A.G., and M.C.N. obtained the resources; all authors contributed writing, review and editing the article.

## Notes

https://4932539.users.earthengine.app/view/sacrconservation

### Competing Interest Statement

The authors have declared no competing interest.

