## AppendixA for "Flood-irrigated agriculture mediates climate-induced wetland scarcity for summering sandhill cranes in western North America"

**Appendix A** - Description of methods used to monitor and evaluate wetland hydroperiod and long-term (1984-2022) wetland trends within core sandhill crane distributions.

Following an approach outlined by Donnelly et al. (2021), wetland surface water hydrology was measured using constrained spectral mixture analysis (SMA, Adams & Gillespie 2006) that allowed proportional estimations of water contained within a continuous 30 x 30 m pixel grid (Halabisky et al. 2016; Jin et al., 2017). This approach accurately accounts for surface water area/extent when detectability is reduced due to interspersion of emergent vegetation, shallow, or turbid water (DeVries et al. 2017); characteristics common to wetlands frequently used by sandhill cranes in the Intermountain West (Drewien & Bizeau 1974). Areas containing cloud, cloud shadows, snow, and ice were masked using the Landsat CFMask band (Foga et al. 2017). All unmasked pixels in Landsat 30 m visible, near-infrared, and short-wave infrared bands were incorporated into the SMA with the exception of Landsat 8 coastal aerosol band.

Training data for SMA were extracted from satellite imagery as spectral endmembers unique to individual images classified. Training site locations represented homogeneous land cover types mapped as water, wetland vegetation, upland, and bare soil. Spectral endmembers for water were collected using image masks generated from 99th percentile normalized difference water index values (McFeeters 1996). Mask extents coincided with large deep water lakes/reservoirs proximal to wetland sites. A similar masking approach was applied to collect wetland vegetation endmembers using normalized difference vegetation indices (Box et al. 1989). Sampling was constrained to sites coincident with seasonally flooded wetlands and were representative of associated plant phenology. Spectral mixture analysis requires minimal training data (Adams & Gillespie 2006), which allowed upland and bare soil endmembers to be generated from a small number of static plots within the study area (n = 2; 0.5-1 $km^2$). Upland plots were associated with homogenous shrublands with low vegetative productivity and high soil exposure. Bare soil plots were coincident with dry lake basins in areas of surface mineral deposits. Plot locations were identified using high-resolution (< 0.5 m) multispectral satellite imagery or field reconnaissance.

Using on-screen interpretation, wetland pixels were classified into functional groups to allow specific ecologic and land-use characteristics to be associated with surface water hydrology. This made it possible to stratify wetland data into meaningful summaries and remove features extraneous to summering sandhill crane habitats (e.g., reservoirs, swedge ponds, large rivers). Because emergent vegetation or high turbidity could partially mask pixel areas covered with water (Donnelly et al. 2019), we considered pixels fully inundated when water was present. Pixels containing <15% surface water were omitted from summaries to minimize the overestimation of surface water area.

A series of procedures were used to minimize potential false water positives, including the use of slope models derived from the Shuttle Radar Topography Mission (SRTM) digital elevation dataset (Jarvis et al. 2008). Areas of high topographic relief were used to mask terrain shadows known to negatively influence water detection when using SMA (DeVries et al. 2017). Additionally, the impervious surfaces band from the U.S. Geological Survey National Land Cover Database (NLCD; Dewitz 2023) was used to remove false surface water positives attributed to urban features (e.g., asphalt surfaces and building shadows). Shadows associated with coniferous forests and woodlands were also removed using the NLCD by selecting associated land cover classes and applying them as masks. Accuracy was estimated to be 93-98% by comparison to previous work and similar methods used by Donnelly et al. (2019) that overlapped a quarter of our study site. This estimate was comparable to similar time-series wetland inundation studies using Landsat data (Jin et al. 2017).

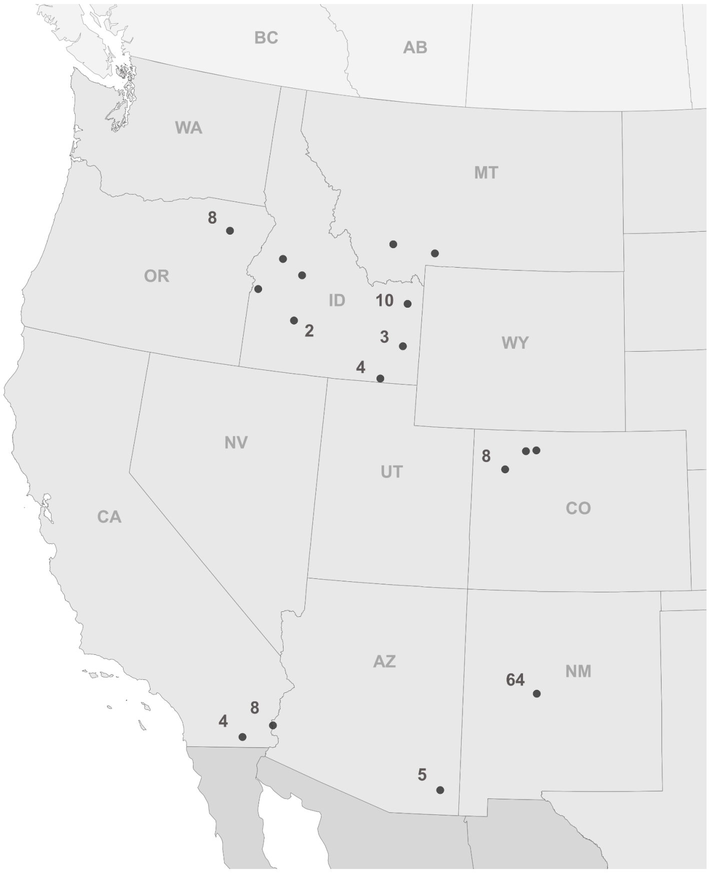


Figure A1. Sandhill crane capture locations (black points). Numbers indicate total GPS band deployed is greater than one.
