## AppendixB for "Flood-irrigated agriculture mediates climate-induced wetland scarcity for summering sandhill cranes in western North America"

Appendix B - Sandhill crane wetland trend and land ownership summaries

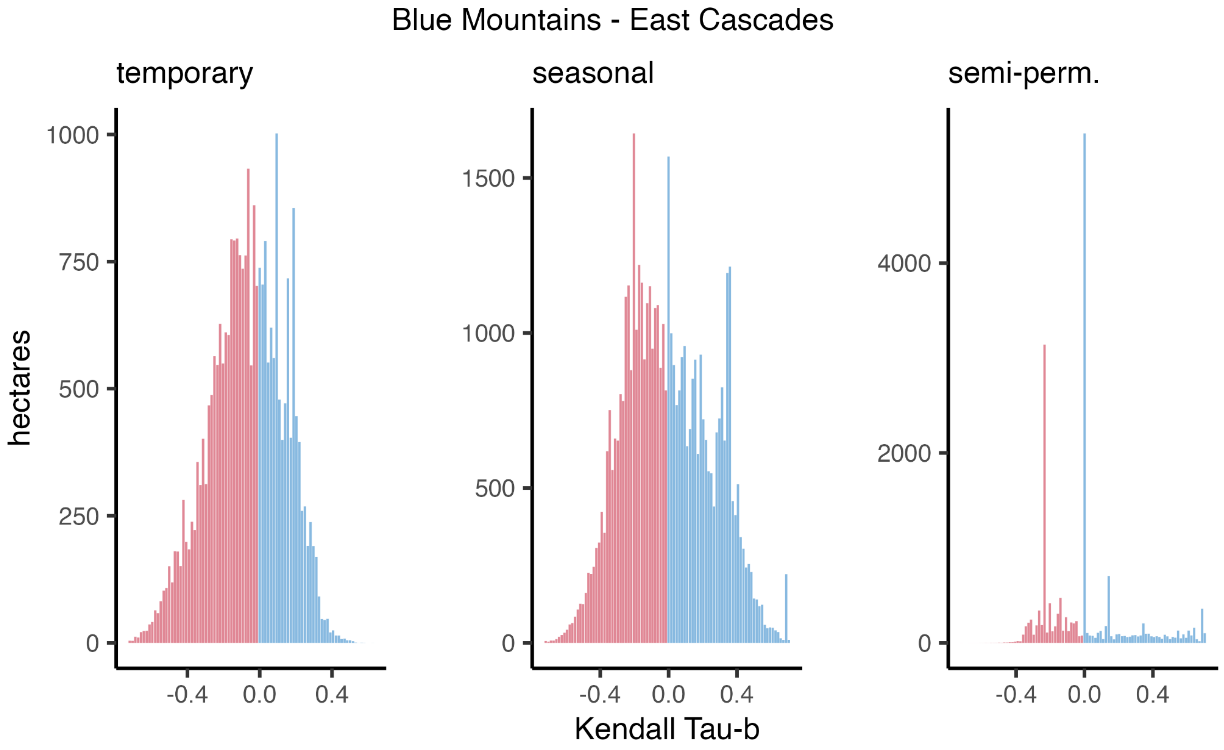


Figure B2. Blue Mountains - East Cascades Kendall's Tau-b rank correlation value distributions by wetland hydroperiod class. Summaries represent all wetlands within sandhill crane core areas. Negative values indicate wetland drying associated with reduced duration or frequency of flooding.

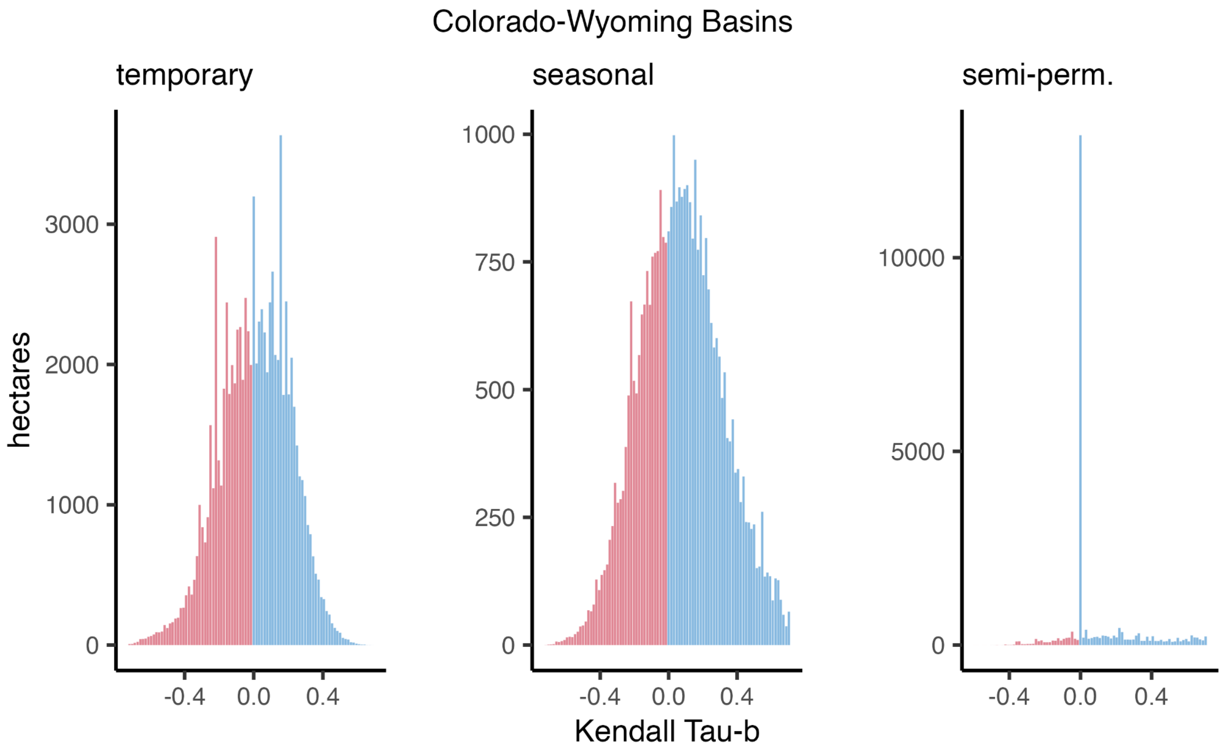


Figure B3. Wyoming - Colorado Bains Kendall's Tau-b rank correlation value distributions by wetland hydroperiod class. Summaries represent all wetlands within sandhill crane core areas. Negative values indicate wetland drying associated with reduced duration or frequency of flooding.

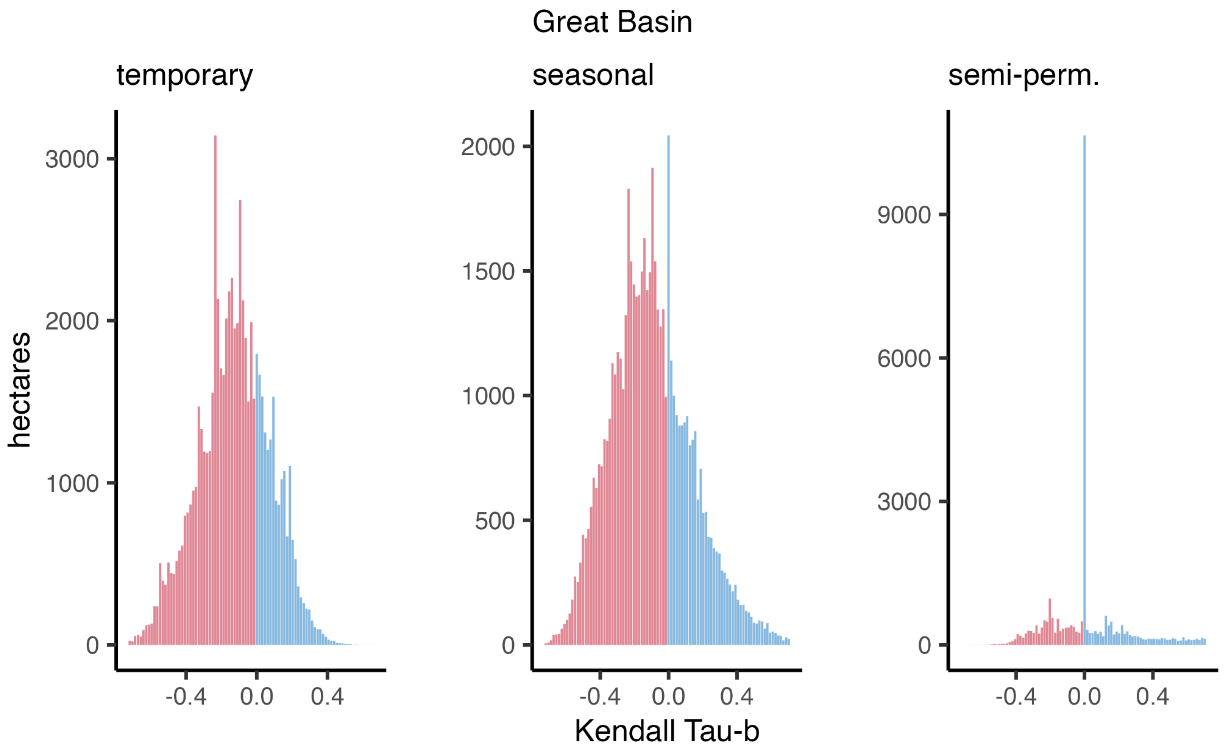


Figure B4. Great Basin Kendall's Tau-b rank correlation value distributions by wetland hydroperiod class. Summaries represent all wetlands within sandhill crane core areas. Negative values indicate wetland drying associated with reduced duration or frequency of flooding.

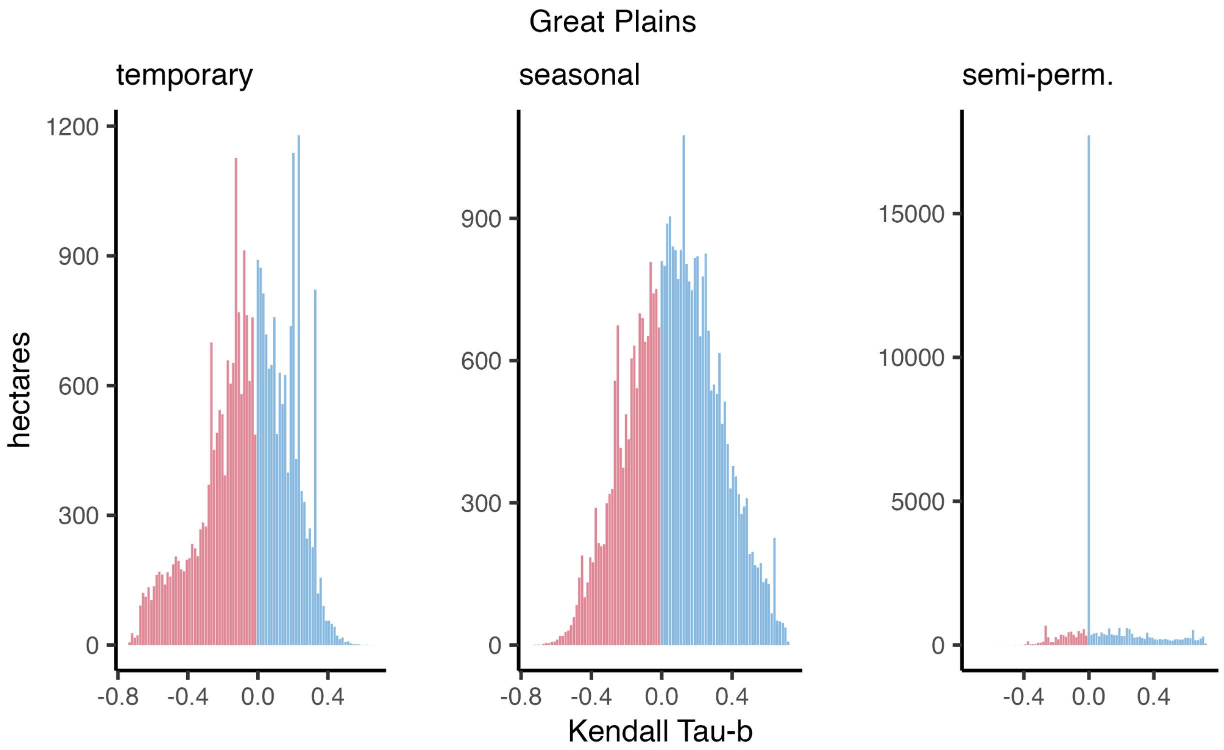


Figure B5. Great Plains Kendall's Tau-b rank correlation value distributions by wetland hydroperiod class. Summaries represent all wetlands within sandhill crane core areas. Negative values indicate wetland drying associated with reduced duration or frequency of flooding.

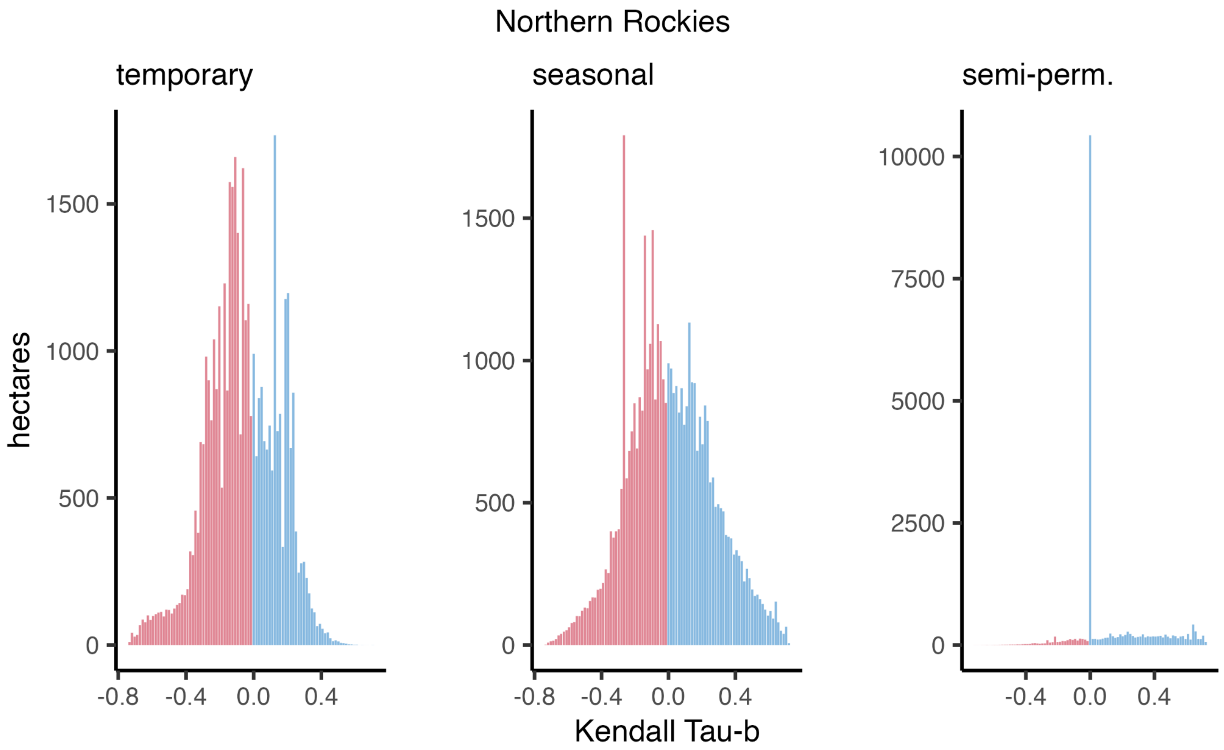


Figure B6. Northern Rockies Kendall's Tau-b rank correlation value distributions by wetland hydroperiod class. Summaries represent all wetlands within sandhill crane core areas. Negative values indicate wetland drying associated with reduced duration or frequency of flooding.

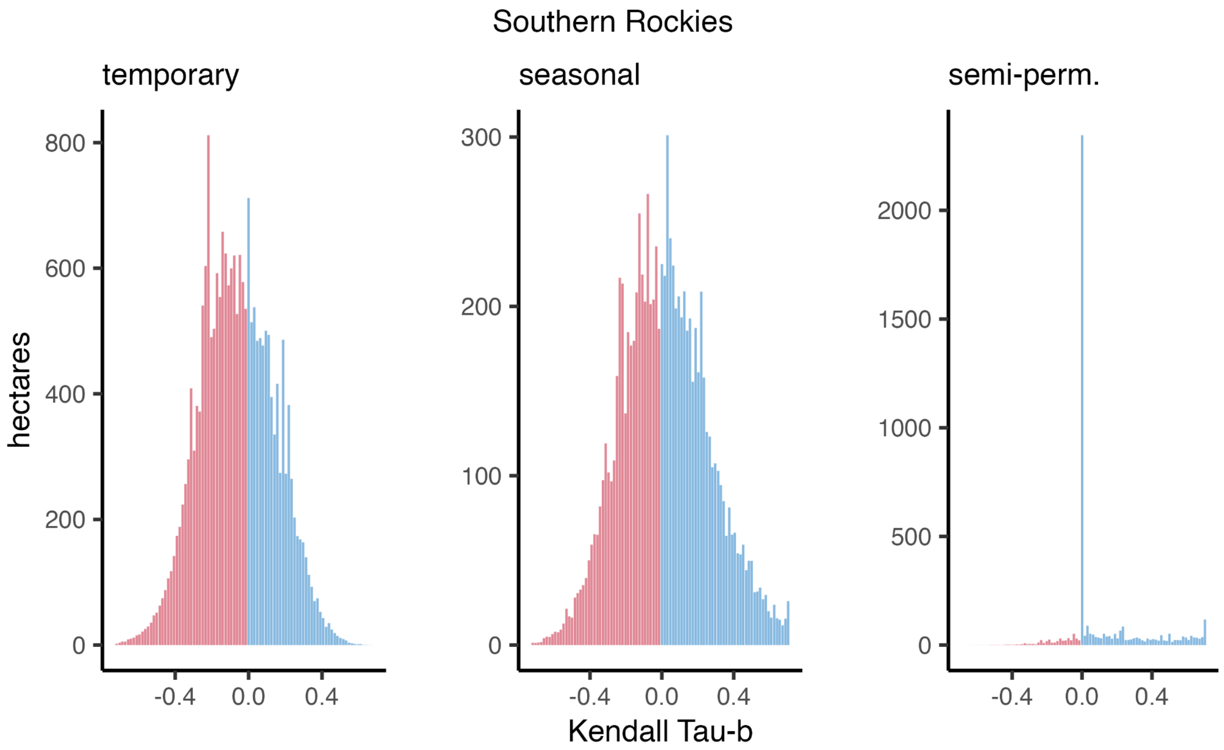


Figure B7. Southern Rockies Kendall's Tau-b rank correlation value distributions by wetland hydroperiod class. Summaries represent all wetlands within sandhill crane core areas. Negative values indicate wetland drying associated with reduced duration or frequency of flooding.

Table B1. Sandhill crane core summering area summarized by public land management agency. Percentages represent proportional abundance within the overall core area. Other public lands class represents an amalgamation of over 15 agencies and municipalities that individually account for ≤ 0.3% of core summering areas.

| Public land management agency | hectares | percentage |
|---|---|---|
| U.S. Forest Service | 57,439 | 4.5% |
| Bureau of Land Management | 34,382 | 2.7% |
| U.S. Fish and Wildlife Service | 31,963 | 2.5% |
| State Wildlife Agencies | 18,704 | 1.5% |
| National Park Service | 10,612 | 0.8% |
| U.S. Bureau of Reclamation | 5,916 | 0.5% |
| Other public lands | 43,268 | 3.4% |

Table B2. Temporary wetlands summarized by public land management agency. Percentages represent proportional abundance within the overall sandhill crane core summering area. Other public lands represent an amalgamation of minor occurrences from over 15 agencies and municipalities.

| Public land management agency | hectares | percentage |
|---|---|---|
| U.S. Fish and Wildlife Service | 22,969 | 3.1% |
| U.S. Forest Service | 16,867 | 2.3% |
| State Wildlife Agencies | 13,074 | 1.8% |
| Bureau of Land Management | 10,580 | 1.4% |
| National Park Service | 3,838 | 0.5% |
| Other public lands | 21,260 | 2.9% |

Table B3. Seasonal wetlands summarized by public land management agency. Percentages represent proportional abundance within the overall sandhill crane core summering area. Other public lands represent an amalgamation of minor occurrences from over 15 agencies and municipalities.

| Public land management agency | hectares | percentage |
|---|---|---|
| U.S. Fish and Wildlife Service | 60,676 | 10.7% |
| U.S. Forest Service | 16,867 | 5.8% |
| State Wildlife Agencies | 29,244 | 5.2% |
| Bureau of Land Management | 20,381 | 3.6% |
| National Park Service | 7,525 | 1.4% |
| Other public lands | 21,911 | 3.9% |

Table B4. Semi-permanent wetlands summarized by public land management agency. Percentages represent proportional abundance within the overall sandhill crane core summering area. Other public lands represent an amalgamation of minor occurrences from over 15 agencies and municipalities.

| Public land management agency | hectares | percentage |
|---|---|---|
| U.S. Fish and Wildlife Service | 42,673 | 12.9% |
| U.S. Forest Service | 8,269 | 2.5% |
| State Wildlife Agencies | 29,365 | 8.9% |
| Bureau of Land Management | 15,576 | 4.7% |
| National Park Service | 2,037 | 0.6% |
| Other public lands | 8,387 | 3.6% |
